# Snapper: a high-sensitive algorithm to detect methylation motifs based on Oxford Nanopore reads

**DOI:** 10.1101/2022.11.15.516621

**Authors:** Dmitry N. Konanov, Vladislav V. Babenko, Aleksandra M. Belova, Arina G. Madan, Daria I. Boldyreva, Oksana E. Glushchenko, Ivan O. Butenko, Dmitry E. Fedorov, Aleksandr I. Manolov, Vassili N. Lazarev, Vadim M. Govorun, Elena N. Ilina

**Affiliations:** Research Institute for Systems Biology and Medicine, Moscow, Russia; Federal Research and Clinical Center of Physical-Chemical Medicine, Federal Medical Biological Agency, Moscow, Russia

## Abstract

The Oxford Nanopore technology has a great potential for the analysis of genome methylation, including full-genome methylome profiling. However, there are certain issues while identifying methylation motif sequences caused by low sensitivity of the currently available motif enrichment algorithms. Here, we present Snapper, a new highly-sensitive approach to extract methylation motif sequences based on a greedy motif selection algorithm. Snapper has shown higher enrichment sensitivity compared with the MEME tool coupled with Tombo or Nanodisco instruments, which was demonstrated on *H. pylori* strain J99 studied earlier using the PacBio technology. In addition, we used Snapper to characterize the total methylome of a new *H*.*pylori* strain A45. The analysis revealed the presence of at least 4 methylation sites that have not been described for *H. pylori* earlier. We experimentally confirmed a new CCAG-specific methyltransferase and indirectly inferred a new CCAAK-specific methyltransferase.

## Introduction

Restriction-modification (R-M) systems are one of essential mechanisms used by bacteria for protection from bacteriophage invasion. Generally, bacterial R-M systems are divided into four global groups (I, II, III, and IV) [1] in dependence of the R-M complex structure, but the common feature of them is the presence of site-specific DNA-binding domains which recognize specific short DNA sequence that should be methylated to prevent the restrictase activity.

The Oxford Nanopore technology (ONT) is a long-read sequencing technology based on measuring electric current across the nanopore [2]. The ONT makes it possible not only to detect the four canonical bases, but also to additionally catch any of their modified forms that can significantly change the sequencing signal [3]. A number of tools aimed at accurate detection of methylated positions have already been developed [4-7]. However, these tools have certain limitations while the inference of methylation site sequences, especially when processing low-represented DNA motifs. Here, we present a novel tool called Snapper that performs high-sensitive detection of methylation motifs based on ONT sequencing data. The tool is verified on *Helicobacter pylori* J99 sequencing data and compared with Tombo and Nanodisco which are the most up-to-date instruments with similar functionality.

We chose *H. pylori* species as the object of interest in this study firstly because it is a unique organism that is shown to bring up to 30 different R-M systems in its genome [8] and has methyltransferases(MTases) capable of methylation adenine as well as cytosine [8]. Secondly, it is a naturally competent organism [9] that can be quite easily modified to confirm new inferred MTases. Third, the *H. pylori* J99 strain has been characterized earlier using the PacBio sequencing technology [10] that allows us to directly compare our results with competing technology.

To demonstrate the applicability of the developed tool, here we characterize the total methylome of a new *H. pylori* strain A45 using Snapper and additionally show the method accuracy on A45 mutants disrupted in genes of methyltransferases with known specificity.

## Results

### Software description

We have developed a novel high-sensitive algorithm for methylation motifs detecting based on Oxford Nanopore sequencing data. The algorithm has been implemented as a command-line tool called Snapper and is available as a pip package to install. To perform non-target methylome analysis, the tool requires native+control DNA samples being sequenced and resquiggled using Tombo. Snapper uses a k-mer approach, with k chosen to be 11 in order to cover all 6-mers that cover one particular base (Figure 1, A) under the assumption that, in general, approximately 6 bases are located in the nanopore simultaneously [11].

**Figure 1.**
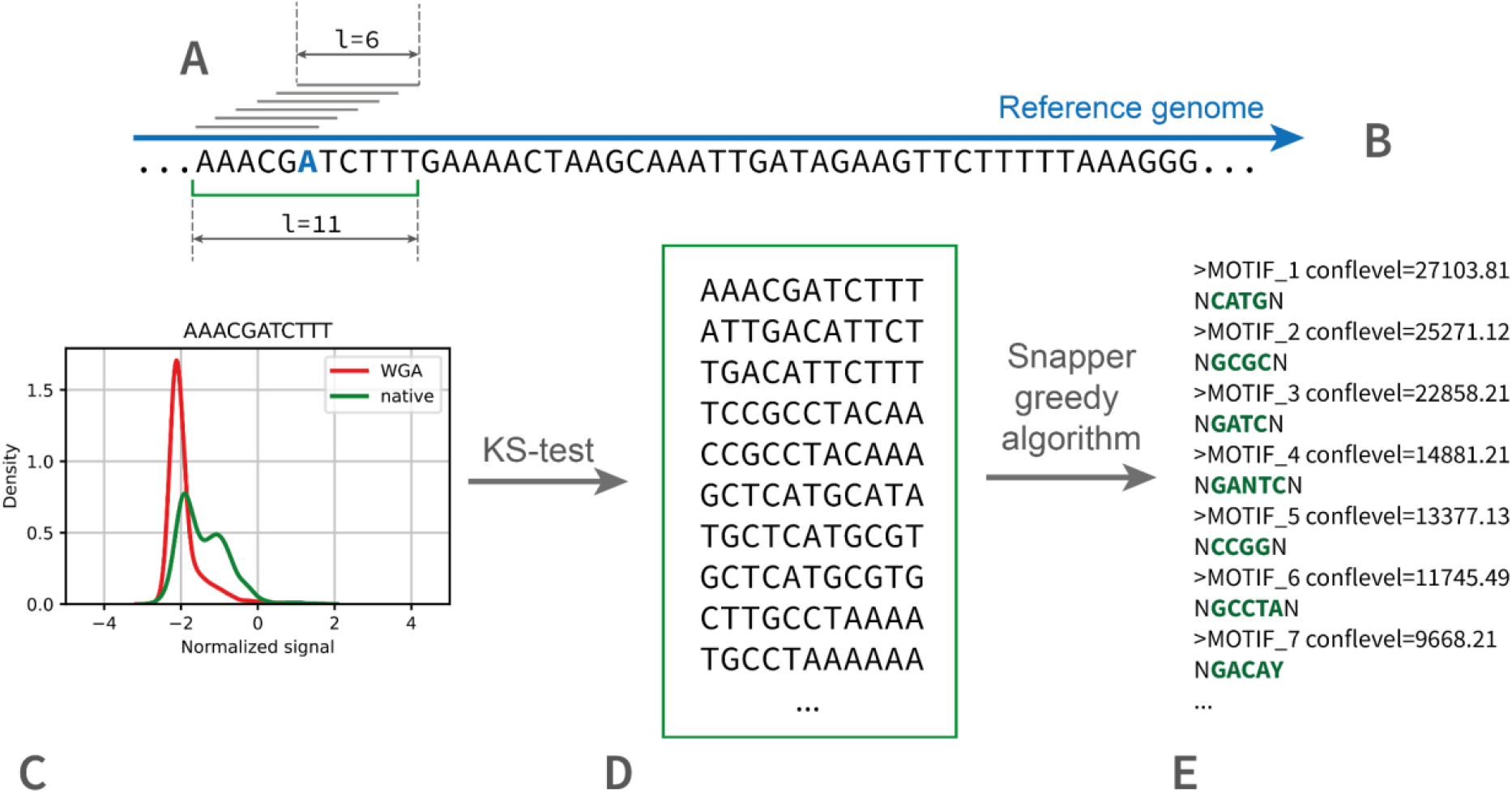
The principal scheme of the Snapper pipeline. In the first stage, for each 11-mer in the reference genome (**A-B**), the algorithm collects normalized signal levels for this 11-mer from multi-fast5 files for both native and WGA samples. Thereafter, for each 11-mer presented in the genome, the algorithm directly compares the collected signal distributions (**C**) using the Kolmogorov-Smirnov test in order to select 11-mers that most likely contain a modified base. The result of the first stage is an exhaustive set of all potentially modified 11-mers (**D**). Next, the greedy motif enrichment algorithm implemented in Snapper iteratively extracts potential methylation motifs and calculates corresponding motif confidence levels (**E**). By default, motifs with confidence level greater than 1000.0 are considered as significant but the authors recommend manually verifying extracted motifs with confidence levels lower than 5000 especially while the observed signal shift is rather weak. The main recommendations on how to interpret the Snapper results are available in the user guide (https://snapper-tutorial.readthedocs.io/en/latest/usercases.html#main-points).

The first stage of the Snapper pipeline is the extraction of nucleotide 11-mers which are most likely to contain a modified base. The comparison of signal distributions (Figure 1, C) is performed using the Kolmogorov-Smirnov test statistics. Next, all the extracted k-mers (Figure 1, D) are merged by a greedy algorithm which generates the minimal set of potential modification motifs which can explain the most part of selected 11-mers (Figure 1, E), under the assumption that all selected 11-mers contain at least one modified base. Another feature of Snapper is a transparent way to estimate a confidence level of extracted motifs using chi-square statistics. It might be useful while manual observation of Snapper results since motifs with quite low confidence level (usually less than 3000) in some cases should be manually adjusted or just removed. The more detailed algorithm explanation is available in the Methods section and Supplementary Materials 9. A short tool’s guideline is available on https://snapper-tutorial.readthedocs.io. All the source code is available on GitHub (https://github.com/DNKonanov/Snapper)

Snapper output folder looks as following:

- “passed_motifs_[strand]_[contigname].fasta” - an exhaustive set of 11-mers which are most likely to contain a methylated base
- “final_motifs_[strand]_[contigname].fasta” - a list of potential methylation motifs, generated by the greedy algorithm and corresponding confidence levels
- “seqs_iter” folder contains fasta files that contain unexplained 11-mers for each greedy algorithm work iteration. The “seqs_iter_1.fasta” file is equivalent to “passed_motifs_*” for this particular contig and strand.
- “plots_[strand]_[contigname] folder contains signal distribution plots for all extracted methylation motifs. These plots can be used to additionally verify the results.

The tool has a few optional parameters. Firstly, the user can tune the threshold value of KS-test -log10(p-value) on the 11-mers selection stage. Secondly, the greedy motif extraction algorithm has two main parameters to tune; the first *confidence* parameter defines the minimal value of the chi-square statistics used for the motif extraction, the second *max_motifs* parameter limits the number of potential motifs extracted. Tuning of KS-test p-value is recommended by the authors only in cases when the mean sequencing coverage of either native or control sample DNA is lower than 20. The tuning of the motif extraction parameters depends on the considered organism. Thus, *Helicobacter pylori* considered in this study has been shown to have up to 20 Type II-III different R-M systems encoded in the genome, so we chose the desired number of motifs to be 20.

In order to verify the approach developed, firstly we performed the total methylome analysis of *Helicobacter pylori* J99 strain which has been characterized earlier using competing long-read sequencing technology PacBio [10]. Additionally, we compared our method with instruments Tombo [4] and Nanodisco [5] which are in our knowledge the most up-to-date tools capable of non-target methylation motifs profiling besides Snapper.

### Total methylome analysis of *H. pylori* J99

The genome of *Helicobacter pylori* J99 (genotype cagA+/vacA s1m1) encodes about 20 R-M systems of Type II and Type III [12]. At least 14 out of them have been shown to be active earlier using the PacBio long-reads sequencing [10]. We performed the full Snapper pipeline and discovered the same set of motifs except TCNNGA (Table 1, ONT+Snapper column. We should note that in Table 1 we demonstrate just the Snapper’s raw output except two false-positive variants. You can see more details in Supplementary Materials 1.) To verify the absence of TCNNGA motif we manually observed a subset of genome contexts that contained this pattern and had a significant signal shift. We found that all such contexts contained other extracted motifs such as GAGG, GATC, TCGA and others. Thus, we concluded that despite the presence of TCNNGA in a number of methylated positions it is not an individual methylation motif in our J99 strain (more information about TCNNGA is available in Supplementary Materials 2). In case of GTSAC and GWCAY motifs we faced a certain ambiguity while motifs merging since GTCAC is a submotif for both these sites. Formally, such cases can be resolved only experimentally and it is the reason why we decided not to merge motifs automatically in the Snapper pipeline.

**Table 1.**
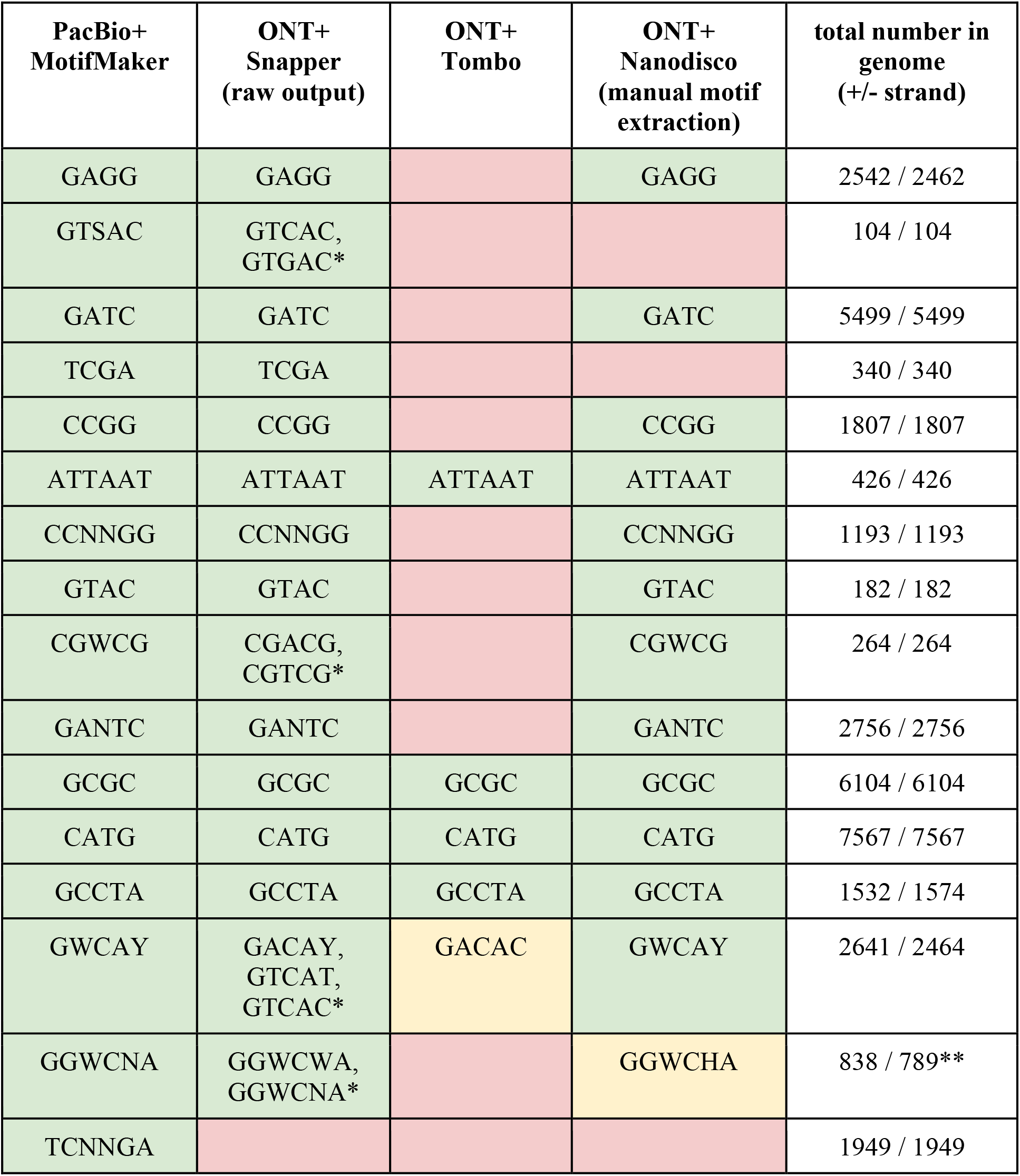
Lists of J99 methylation motifs obtained using different methylation motif detection approaches. It should be noted here, that the column entitled Snapper contains just the raw Snapper output excluding few false-positive motifs that were removed after manual results observation. The raw Snapper output and the verification process of this particular case are available in Supplementary Materials 1. The PacBio results for *H*.*pylori* J99 were taken from the J.Krebes at. all’s work [10] directly (with few changes since the authors considered not the wild J99 strain but J99-3 mutant). For Tombo the best results which we managed to obtain with MEME are shown. For Nanodisco, the manually curated results are shown since the results generated in the automated mode were drastically less accurate. * These motifs were extracted as different because the algorithm is designed to be quite cautious in motifs merging to prevent occasional motif collisions. ** For GGWCNA variant

Additionally, we analyzed *H. pylori* J99 ONT reads using Tombo and Nanodisco tools which are mainly designed at methylated position profiling but have their own motif enrichment functionality as well. Tombo uses the default MEME motif enrichment algorithm [13] to extract motifs, so it has demonstrated rather low sensitivity and has inferred only five motifs highly-represented in the genome (Table 1, ONT+Tombo column). Nanodisco also performs motif enrichment with MEME but uses an iterative approach that allows to detect more methylation sites but requires manual control on each motif extraction iteration because as we found in the fully automated mode Nanodisco is strongly prone to excessive motif collisions and even the most-represented motifs such as CATG and GCGC are not correctly discovered automatically. Generally, manually curated Nanodisco results are quite close to the automated Snapper output and the PacBio results except GTSAC and TCGA motifs (Table 1, ONT+Nanodisco column). Both these motifs are rather rare in the J99 genome, so the MEME algorithm did not identify them being enriched even using an iterative approach since Nanodisco uses the default MEME objective function without negative background (according to the Nanodisco source code). In Snapper, a set of all possible 11-mers presented in the considered genome is used as a negative background.

### Total methylome analysis of *H. pylori* A45 and mutants

The *H. pylori* A45 (genotype cagA-/vacA s2m2) clinical isolate was obtained previously from a gastric mucosa biopsy sample from a patient with a gastric carcinoma [14]. In addition, the derivatives of the A45 strain of *H. pylori*, defective in genes methyltransferases M.HpyAI (*hpy* mutant), M.HpyAIII (*hp91/92* mutant), M.HpyAIV (*hp1352* mutant) were obtained [14].

The analysis of *H. pylori* A45 has revealed presence of 15 methylation sites belonging to R-M systems of types II and III: ATTAAT, GTNNAC, GGRGA, CCATC, CATG, CCAG, GCGC, GANTC, GATC, GGCC, GAAC, TGCA, TCGA, TCNGA, TCNNGA (Table 2, wild type column).

**Table 2.** Methylation motifs detected in the wild A45 and four mutants. Interestingly, in the *hpy* and *hp91/92* mutants an additional CCAAK methylation motif can be observed.

| wild type | total number in genome (+/- strand) | *hpy* | *hp1352* | *hp91/92* | *hp944* |
| --- | --- | --- | --- | --- | --- |
| ATTAAT | 441 / 441 | ATTAAT | ATTAAT | ATTAAT | ATTAAT |
| GTNNAC | 325 / 325 | GTNNAC | GTNNAC | GTNNAC | GTNNAC |
| GGRGA | 1768 / 1790 | GGRGA | GGRGA | GGRGA | GGRGA |
| CCATC | 1163 / 1115 | CCATC | CCATC | CCATC | CCATC |
| CATG | 7246 / 7246 | - | CATG | CATG | CATG |
| CCAG | 2173 / 2271 | CCAG | CCAG | CCAG | - |
| GCGC | 5982 / 5982 | GCGC | GCGC | GCGC | GCGC |
| GANTC | 2625 / 2625 | GANTC | - | GANTC | GANTC |
| GATC | 5113 / 5113 | GATC | GATC | - | GATC |
| GGCC | 1534 / 1534 | GGCC | GGCC | GGCC | GGCC |
| GAAC | 2760 / 2700 | GAAC | GAAC | GAAC | GAAC |
| TGCA | 5770 / 5770 | TGCA | TGCA | TGCA | TGCA |
| TCGA | 290 / 290 | TCGA | TCGA | TCGA | TCGA |
| TCNGA | 1224 / 1224 | TCNGA | TCNGA | TCNGA | TCNGA |
| TCNNGA | 1926 / 1926 | TCNNGA | TCNNGA | TCNNGA | TCNNGA |
| - | 3403 / 3267 | CCAAK | - | CCAAK | - |

Interestingly, opposite to the J99 analysis, in this case all the extracted motifs had quite high confidence levels greater than 3000 (Supplementary Materials 3). Despite that, we manually observed signal distributions for all suggested sites and their closest genome context to ensure that all motifs had been extracted correctly.

To identify the MTases responsible for methylation of new sites CCAG, GGRGA, and GAAC not described earlier for *H. pylori*, which were presented in *H*.*pylori* A45 but not presented in J99, all candidate MTase genes were extracted from both genomes and used to construct a phylogenetic tree based on protein sequences similarity. Two proteins that were presented only in A45 strain and did not have orthologous genes in J99 were considered as potentially new MTases. Two new mutants (*hp8* and *hp944*) disrupted in corresponding genes *hp0008* and *hp0944* were obtained during this work, their native DNA was analyzed using ONT and the sequencing data were processed using Snapper. *hp8* did not have any signal changes, but in *hp944* we observed a successful deactivation of CCAG-specific MTase (Table 2, *hp944* column). As we found later, the negative result for *hp8* was caused by a nonsense mutation in the MTase gene.

Here, to estimate the method specificity, in addition to the wild type we analyzed three mutants of *H. pylori* A45 disrupted in three different genes encoding MTases with known specificity (*hpy, hp1352* and *hp91/92* mutants disrupted in the genes of MTases specific to CATG, GANTC and GATC motifs respectively). Here, we used as a control not the WGA sample but the native A45 to check out how the algorithm works with a small number of motifs that differ by their signal level. We expected the algorithm to extract only one motif for each mutant, but two mutants had an additional motif that seemed to be modified. Firstly, in all three mutants Snapper successfully detected the absence of methylation of the corresponding motifs (Table 2, *hpy, hp1352*, and *hp91/92* columns, and Supplementary Figure 3). Secondly, the CCAAK site was detected as methylated in *hpy* and *hp91/92* mutants while the native sample, *hp1352*, and *hp944* had CCAAK signal distributions identical to the WGA sample (Figure 2, A).

**Figure 2.**
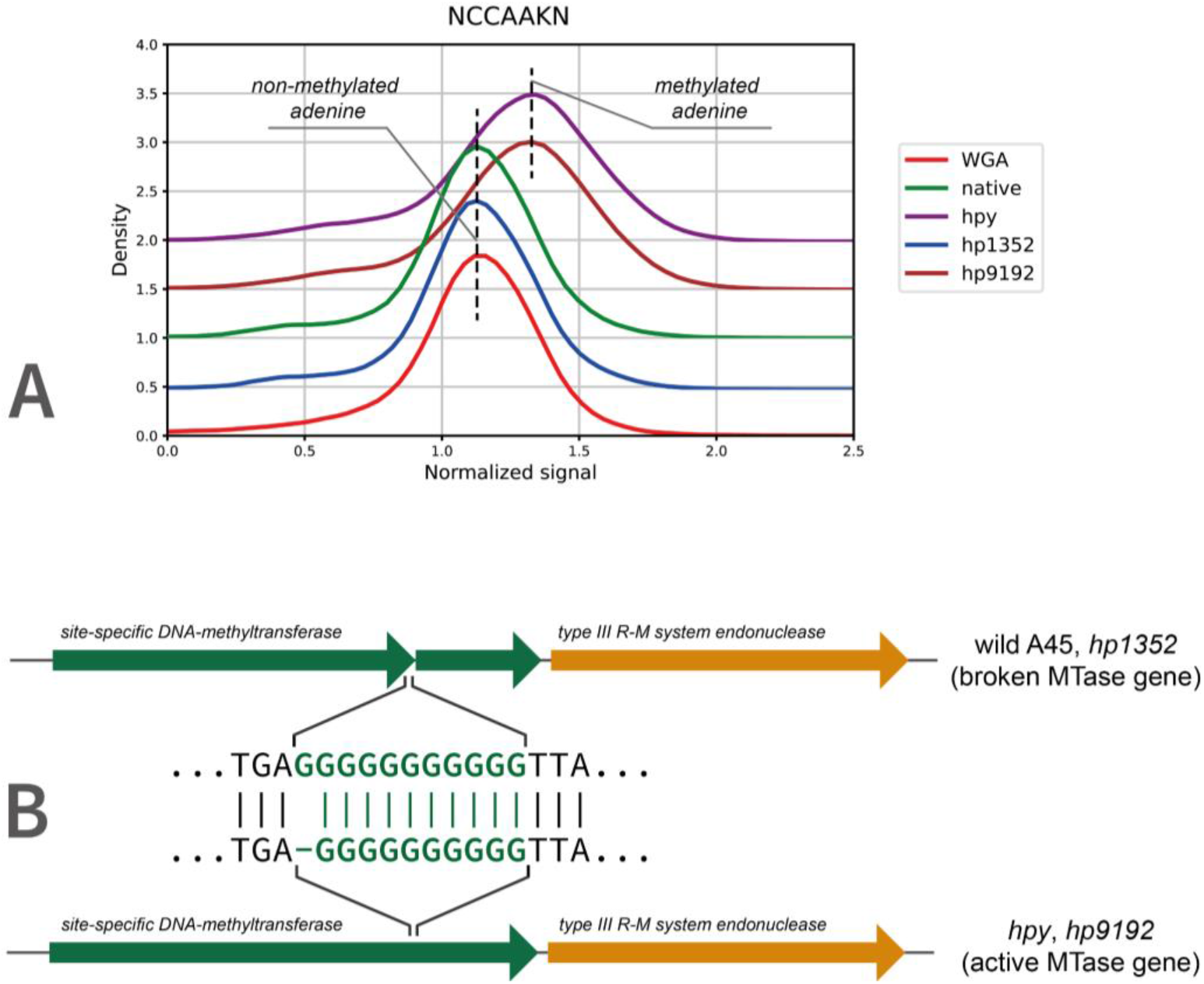
**A)** Normalized signal distributions for the CCAAK methylation site. *hp91/92* (brown) and *hpy* (purple) mutants have a visible signal shift in comparison with the WGA sample (red) while the wild A45 (green) and the *hp1352* (dark blue) mutant have signal distributions identical to the WGA sample (red). **B)** Wild A45 and the *hp1352* mutant have CCAAK-specific MTase gene broken because of a frame-shift insertion in a homopolymeric region in the gene. The reverse frame-shift mutation in the MTase gene restores the Type III R-M system activity in *hpy* and *hp91/92*.

To identify the MTase responsible for modification of CCAAK, we used non-target proteomics data that were obtained earlier for the wild A45, and *hpy, hp1352*, and *hp91/92* mutants. Five proteins that were significantly overrepresented (LFC > 5) in *hpy* and *hp91/92* mutants in comparison to the wild type and *hp1352* were found and manually annotated using BLASTP (Supplementary Tables 10-11). Proteins encoded by *hp0008* and *hp0009* genes were annotated as two fragments of one MTase gene, and gene *hp0010* was annotated as a Type III R-M system endonuclease, so, these three genes encode a full Type III R-M system but with a broken MTase gene. The appearance of the CCAAK methylation site in *hpy* and *hp91/92* was caused by a reverse frame-shift mutation in a homopolymeric region in *hp_0008* gene, which was confirmed by short Illumina reads (Figure 2, B; Supplementary Figure 4) (H. pylori A45 wild type: GCA_000333835.2, *hpy*: GCA_013122115.1, *hp1352*: GCA_013122055.1, *hp91/92*: GCA_013122035.1.).

## Discussion

Oxford Nanopore sequencing has a great potential for the analysis of epigenetic modifications of the four canonical bases. Unfortunately, currently existing software aimed at methylation sites detection has certain drawbacks. Thus, both Tombo and Nanodisco provide high-quality profiling of methylated positions in a genome but the main problem here is to accurately generate a set of potential motifs which could explain the all methylated bases. Technically, it is a classical motif enrichment task, but classical approaches turned out to be not sensitive enough while analysis of highly-methylated genomes such as *H. pylori* genome. Thus, we found that the most widely used tool suit MEME is capable of detecting only highly-represented motifs. An iterative approach implemented in Nanodisco is more sensitive but requires a lot of manual work while extracting motifs that limits its applicability when more than few genomes are studied.

We have developed a new greedy algorithm that is aimed at high-sensitive position-specific motif enrichment. This algorithm has been implemented into command-line tool Snapper. Here, we have shown that, firstly, as a fully automated pipeline Snapper outperforms both Tombo and Nanodisco coupled with the MEME tool and, secondly, Snapper is capable of detecting methylation site sequences with sensitivity equivalent to the PacBio technology and is slightly more sensitive than Nanodisco in the manual mode. In contrast to Nanodisco, Snapper provides a fully-automated motif extraction algorithm and requires manual results verification only while adjustment of motifs with a low confidence level.

In this initial release, we focused only on the short methylation sites detection. Formally, the developed motif enrichment algorithm can be used for identifying longer motif sequences but in this case it would be drastically more time-consuming and require specific search heuristics used that would complicate the algorithm core logic and results transparency. A feature we should note is an intended algorithm cautiousness in motifs merging to avoid occasional motif collisions. Thus, the J99 analysis has shown the presence of motifs GTCAT, GTCAC, GACAY, and GTGAC as methylated. Actually, these submotifs are explained by GWCAY and GTSAC methylation sites but GWCAY+GTGAC variant is formally possible as well. Such cases can be resolved only experimentally so we decided not to merge controversial motifs in the Snapper motif extraction algorithm.

Since Snapper aims not at the methylated positions calling but methylation motif sequences, it is relatively less demanding on the input data compared with Nanodisco or Tombo. The genome coverage required for the analysis depends on the object of interest. Thus, while analyzing organisms that have very few different non-overlapped methylation sites, the method can work well even with a mean genome coverage of 15-20x. On the other hand, the more methylation sites in the genome and the more the chance of motifs collision, the deeper coverage is required for the analysis. The authors suggest that in most cases coverage of 80x-100x is absolutely enough for high-accurate methylation motifs detection.

Using Snapper, we fully characterized the total methylome of a new *Helicobacter pylori* strain A45. In this strain, we found three methylation sites that have not been described earlier for *H*.*pylori* (GGRGA, GAAC, and CCAG) and managed to experimentally confirm the methyltransferase specific to CCAG. In addition, during the experiment we observed a frame-shift based phase variation in the gene encoding a new MTase specific to CCAAK methylation site. Thus, we did not observe this site being methylated in the wild A45 culture but it appeared after the disruption in CATG-specific MTase or GATC-specific MTase. In both cases it was caused by the same reverse frame-shift mutation in a homopolymeric region in the gene encoding corresponding MTase. Such a behavior is quite typical for *H. pylori* and is often used to regulate activity of particular MTase genes [15].

Therefore, we tested Snapper in two different use cases. The first was the total methylome analysis of *H. pylori* strains, when we expected to see a lot of motifs extracted. Here, the method turned out to be sensitive enough to extract all methylation sites detected using PacBio for the same bacteria strain. The second case was the analysis of mutants disrupted in one specific MTase gene, when we expected to see only one motif to be extracted. For each mutant, the algorithm successfully identified the deactivated methylation site and additionally detected one extra site in two mutants which was confirmed using genome assembly and proteomics data, so we can conclude that Snapper is suitable for the analysis of the organisms with a low number of active R-M systems as well. An additional advantage we should note is that ONT is capable of discriminating non-methylated cytosine and 5mC directly (ACGT and GCGC methylation sites in J99) while PacBio requires specific technics such as Tet1 mediated 5mC-conversion to be used that complicates an experiment and might negatively influence the method sensitivity.

## Methods

### Proteomics

#### LC-MS analysis

Liquid chromatographic separation was performed on a reverse phase column (15 cm × 75 μm i.d., Agilent Technologies, USA) packed with Zorbax 300SB-C18 resin (particle size – 3 um, pore diameter – 100 A) that was coupled to a Q-Exactive HF hybrid quadrupole-Orbitrap mass spectrometer (Thermo Fisher Scientific, Germany) via a nanoelectrospray source Nanospray Flex (Thermo Fisher Scientific, Germany). Source voltage was set at 2200 V and capillary temperature at 325°C.

Each sample was introduced through EASY-nLC (Thermo, USA) chromatography system in trap-elute configuration (trap column was a 2 cm × 75 μm i.d. Acclaim PepMap column by Dionex, USA, with C18 resin with 3 um-particles with 100 A pores). Samples were introduced onto trap-column with 10 uL of solvent A (0.1% v/v formic acid) at constant pressure 500 bar. Peptides were eluted with a gradient of0 5 to 50 % (v/v) of solvent B (0.1% v/v formic acid, 79.9% v/v acetonitrile) across 60 minutes at flowrate of 500 nl/min in 3 linear steps (10 minutes to 10% B, 25 min to 20% B, 15 min to 35% B, 10 min to 50% B). After each elution system and columns were washed with 100% of solvent B for 10 minutes and regenerated with 5% of solvent B for 20 minutes.

The mass-spectrometer was operated in positive mode in a data-dependent experiment with survey scans acquired at a resolution of 120,000 at m/z 400 within m/z range of 200-1600 with automatic gain control set for 3×106 and maximum injection time of 32 ms. As many as 20 of the most abundant precursor ions with a charge +2 and above from the survey scan were selected for HCD fragmentation. The normalized collision energy was 27. MS2 spectra were acquired at resolution of 7500 at m/z 400, automatic gain control was set for 2×105 and maximum injection time for 32 ms. After fragmentation ions were dynamically excluded from consideration for 45 s.

#### Quantification

Proteins were relatively quantified using the MaxQuant software version 1.6.10.43. Raw files were searched with an integrated Andromeda search engine against the core peptide database. that consisted of peptides that could be produced by strictly one protein. Trypsin/P was specified as the cleavage enzyme, and two missed cleavages were permissible, minimum peptide length was 7. The FDR was set to 0.01 for peptide spectral matches.

Protein abundance was estimated as a total intensity of its 3 most intense peptides. Difference in each protein’s abundance was considered significant if it allowed the FDR to be below 0.05, in addition.

### Sequencing

#### Oxford Nanopore

The DNA was isolated by using the Wizard DNA extraction kit (Promega Corporation, USA) and size selected with optimized solid phase reversible immobilization (SPRI) beads. The DNA concentration and quality were determined on a Qubit 4 Fluorometer and Nanodrop ND-1000 (Thermo Fisher Scientific). The long reads were generated with MinION sequencing (Oxford Nanopore Technologies, UK). The sequencing libraries were prepared using the ligation sequencing kit SQK-LSK109, native barcoding expansion kit EXP-NBD104 and run in a FLO-MIN106 flow cell. Reads were basecalled using Guppy v3.6.1. using default parameters (guppy_basecaller).

#### Illumina

A NEBNext Ultra DNA library prep kit (New England Biolabs, USA) was used to prepare fragment libraries for genome sequencing. Sequencing was performed on the HiSeq 2500 System (Illumina, USA) HiSeq Rapid SBS Kit V2 using a 2 × 250 bp run configuration.

De novo assembly was performed by hybrid assembler Unicycler (v0.4.8) [16] using default parameters. Identification of the protein-coding sequences and primary annotation were performed using PROKKA v1.14.6 [17]. The draft genomes are available in GenBank with the following accession numbers: *H. pylori* A45 wild type: GCA_000333835.2, *hpy*: GCA_013122115.1, *hp1352*: GCA_013122055.1, *hp91/92*: GCA_013122035.1.

#### Whole-genome amplification

The Qiagen REPLI-g Single Cell Kit was used to perform whole-genome amplification to exclude epigenetically modified bases. The method produced micrograms of DNA from 10 ng of input genomic DNA, following the manufacturer’s guidelines and 8 h of amplification time at 30 °C followed by deactivation at 65 °C for 3 min.

### Gene knockout

The *H. pylori* A45 clinical isolate was obtained previously from a gastric mucosa biopsy sample from a patient with a gastric carcinoma [14]. This isolate was used to obtain *hpy, hp91/92, hp135*2, *hp8, hp944* derivatives defective in methyltransferase genes (Supplementary Table 8). A targeted inactivation procedure was performed using the gene knockout method.

### Bacterial strains and culture manipulation

#### Escherichia coli

strain Top10 required for plasmid vectors assembly and production was cultivated at 37°C on solid Luria–Bertani medium or in liquid Luria–Bertani medium with aeration (150 rpm). The transformation procedure was carried out using the «heat shock» transformation protocol (https://international.neb.com/protocols/2012/05/21/transformation-protocol). Chloramphenicol (8 μg/mL, Panreac, Spain) and kanamycin (15 μg/mL, Sigma, USA) were added to the medium for selection.

#### Helicobacter pylori

strains were cultivated for 20–48 h at 37°C under microaerophilic conditions on Columbian agar solid medium (Becton Dickinson, USA) supplemented with 10% donor horse serum (PAA Labs, Austria). Chloramphenicol (8 μg/mL) and kanamycin (15 μg/mL) were added for the selection and passage of resistant mutant strains. Amplicon cell transformation was carried out as described in [18]. Cells were cultivated for 24 h at 37°C under microaerophilic conditions on solid medium. The cell suspension (10^9^ CFU) was moved on a new plate and cultivated for 4-5 h for undergrowth. Thereafter sterile water suspension, containing 500 ng DNA fragment was applied dropwise onto the solid medium surface containing undergrowth cells. The dishes were then left agar side up in the CO_2_ incubator for 17–20 h. The next day cells were subsequently transferred to a selective medium containing corresponding antibiotics. Clones found on plates after 4-5 days were verified to be *H. pylori* and to contain the specific insertions by PCR using primer sets listed in Supplementary Table 7. For any consequent assay all *H. pylori* strains were harvested from the plates after 2 days growth. Cells were washed with HBSS buffer, pH 7.5 (Hank’s Balanced Salt Solution, Thermo Fisher, United States) and precipitated by centrifugation at 3000 × *g* for 10 min for genomic DNA preparation as described earlier.

### DNA manipulation

Vectors and mutant strains used in this study are listed in Supplementary Supplementary Table 8. All standard methods of DNA manipulation, such as plasmid isolation by alkaline lysis, restriction endonuclease digestion and ligation, were performed according to the protocols of Sambrook et al. [19]. *H. pylori* genomic DNA was prepared using the diaGene kit for DNA extraction from cell cultures (diaGene, Russia). Single DNA fragments or PCR amplification products for cloning or sequencing purposes were purified from agarose gels using Cleanup Standard Kit (Evrogen, Russia). DNA restriction enzymes were obtained from Fermentas (Thermo Fisher Scientific, USA) and were used according to the directions of the manufacturers.

#### Primer designs

All of the primers used in this study for PCR were designed by using Vector NTI Software and presented in Supplementary Table 7.

#### *H. pylori* mutant strains construction

*H. pylori* mutant strains *hpy, hp91/92, hp135*2 were obtained earlier [14]. A cassette containing kanamycin resistance gene (*aphA-3*) as a selectable marker was inserted into the target region of the genome. For a detailed description of the plasmids used to inactivate the genes involved in restriction-modification systems, see Supplementary Materials Table 9. Validation of the correct insertion of the cassette into the *H. pylori* genome was confirmed by PCR-analysis and sequencing of the obtained strains. Target inactivation - genes encoding methyltransferases - was confirmed by restriction analysis of genomic DNA of the obtained strains using methyl-sensitive restriction endonucleases: Mbol (GATC), Hin1ll (CATG), Hinfl (GANTC).

*H. pylori* mutant strains *hp8, hp944* were obtained during this work by homologous recombination. For construction *hp944* mutant strain (A45 strain disrupted in the *hp0944* gene) the amplicon, consisting of chloramphenicol resistance *catGC* cassette flanked by two fragments of the *hp0944* of 363 bp and 456 bp, was obtained by two-step PCR amplification. For construction *hp8* mutant strain (A45 strain disrupted in the *hp0008* gene) the amplicon, consisting of kanamycin resistance gene under chloramphenicol promoter region flanked by two fragments of the *hp0008* gene of 325 bp and 463 bp, was obtained by two-step PCR amplification. All *H. pylori* flanking regions were amplified from genomic DNA of A45 strain using Tersus polymerase kit (Evrogen, Russia) according to the directions of the manufacturers. All corresponding primer sets listed in Supplementary Table 7. Schematic map of the primer location and amplicon structure used for *hp0008* and *hp0944* gene disruption in *H. pylori* A45 strain illustrated by Supplementary Figure 5 in Supplementary Materials.

### Methylation analysis

#### Tombo

Raw signals from the fast5 files obtained for both native and WGS samples were mapped to corresponding reference genome positions using ‘tombo resquiggle’ command [4]. Next, modified positions were extracted using the ‘model_sample_compare’ tombo mode. The resulting positions were used to construct all *k*-mers that are likely to bring a methylation base, with *k* chosen to be 12. These k-meric nucleotide contexts were processed by MEME [13] with algorithm parameters recommended in the Tombo documentation. The motifs that were significantly overrepresented in the selected contexts were considered as methylation sites.

#### Nanodisco

For each strain, both sample and control fast5 files and the assembly file were processed according to the standard Nanodisco (v.1.0.3) pipeline described in the documentation (https://nanodisco.readthedocs.io/en/latest/overview.html). While running motif extraction stage (‘nanodisco motif’), we manually controlled each motif inference iteration observing the pdf files generated by the tool since the automated mode turned out to be prone to occasional motif collisions.

#### Snapper

Single-fast5 files obtained with ‘tombo resquiggle’ were transformed to the multi-fast5 format to provide faster access to the sequencing raw signal data. These multi-fast5 files were processed using the python h5py library in order to collect signals for all k-mers presented in the considered genome for each strand independently. K was chosen to be 11 in order to cover all 6-mers that cover an individual base (Figure 1A). Thus, at first the algorithm generates a hash-table, where all presented 11-mers are used as keys and corresponding normalized signal levels as values.

Next, for each 11-mer in the hash-table, the Kolmogorov-Smirnov test is performed to compare signal level distributions between native and WGA samples (or, more generally, between any two samples passed as the input). Before the testing, the algorithm performs an artificial balancing of the sample sizes by random sampling (without replacement) of N values from each considered signal distribution to ensure the uniformity of the 11-mers coverage (N was chosen to be 200 but can be changed by the user). This balancing procedure is applied to make possible direct comparison of statistics values between k-mers which are differently presented in the reference genome and to use the same statistics thresholds to infer modified positions along the genome. As a result, the algorithm returns the list of 11-mers that are most likely to contain a modified base. In the further text, this list will be named *seq_set*.

The next step differs from the classical motif enrichment approaches like the tools included in the MEME Suit. We have developed a highly-sensitive greedy algorithm that is aimed to generate the minimal set of short supermotifs of length 4-8 that can explain most part of sequences in the *seq_set*, under the assumption that all sequences in the *seq_set* contain at least one modified base. The principal idea of the algorithm is iterative search for the most over-represented motif in the *seq_set* in comparison with a reference genome, extraction it as a potential modification motif, following procedure of motif adjustment which additionally ensures the correctness of the extracted motif, and filtering the *seq_set* list by removing all sequences that contain the extracted motif variant. It should be noted here, that extracted motifs are position-specific, so the same motif can be extracted twice or more from different positions in 11-mer. After the *seq_set* filtering, the search and extraction procedure repeats. The termination of the algorithm work is determined by the input parameters *confidence* and *max_motifs*, where the *confidence* parameter means the minimal value of chi-square test statistic (the default value is 1000.0), and the *max_motifs* parameter means the maximum desired number of extracted modification motifs (the default value is 20). As a result, the algorithm returns a sorted list of all potential motifs with corresponding confidence levels (we observed that typical confidence values for known methylation motifs are higher than 3000). A more formal explanation of the motif enrichment algorithm is available in Supplementary Materials 12.

## Software used

Snapper is implemented using Python 3.7. FAST5-files are processed with the h5py library. Sequence processing is performed using the Biopython Python library. The numpy library and regular expressions were used for fast motifs counting and matching. Signal distribution plots are created using matplotlib.

## Data availability

The full data set including single-fast5, multi-fast5, fastq and sequencing summary files had a total size more than 1Tb. Unfortunately, at the current moment there are no suitable online resources for uploading fast5 datasets. Instead, we provide a small example dataset in order to demonstrate the tool capabilities. The demo-dataset includes 10+10 multi-fast5 files containing 40000 long reads for the *hpy* mutant and 40000 reads for the wild A45 as a control (http://download.ripcm.com/snapper_test/). All genome assemblies are available in GenBank (*H. pylori* A45 wild type: GCA_000333835.2, *hpy*: GCA_013122115.1, *hp1352*: GCA_013122055.1, *hp91/92*: GCA_013122035.1).

## Code availability

All the source code is available on Github (https://github.com/DNKonanov/Snapper).

## Competing interests

The authors have no competing interests.

Acknowledgements

We thank the Center for Precision Genome Editing and Genetic Technologies for Biomedicine, Federal Research and Clinical Center of Physical–Chemical Medicine of the Federal Medical Biological Agency for providing computational resources for this project. *H. pylori* J99 strain (№ 700824, cagA+/vacA s1m1 genotype) was kindly provided by prof. M. Achtman (Max Planck Institute for Infection Biology, Germany). *H. pylori* A45 and *hpy, hp1352*, and *hp91/92* mutants were kindly provided by Dr. K.T. Momynaliev.

## Authors’ contributions

D.N.K. and V.V.B. led the project, wrote the manuscript text and prepared all the figures and supplementary data, D.N.K. designed and implemented the main algorithms and performed *H*.*pylori* A45 and J99 methylome analysis, A.M.B. wrote the manuscript text, realized cell culture procedures and prepared *H. pylori hp8* and *hp944* mutant strains, A.G.M. ran Nanodisco and helped with multiprocessing implementation, O.E.G. ran Tombo and helped with methylation analysis, I.O.B. performed proteomics experiments, D.E.F. and A.I.M. processed the proteomics data, A.I.M. and V.V.B. carried out genomic data analysis, V.V.B. and D.I.B. performed genome sequencing, V.N.L., V.M.G. and E.N.I. administered the project and edited the manuscript.

