## Supplementary Materials for "Snapper: a high-sensitive algorithm to detect methylation motifs based on Oxford Nanopore reads"

### 1. J99 extracted motifs

**Supplementary Table 1.** Snapper output for *H. pylori* J99

| MOTIF | conflevel | effsize | comment |
|---|---|---|---|
| NCATGN | 30725.1 | 1.304 | |
| NGCGCN | 31789.6 | 0.85 | |
| NGATCN | 28787.6 | 0.36 | extremely high confidence level |
| NGANTCN | 18370.6 | 0.4 | high confidence level |
| NCCGGN | 16504.4 | 0.98 | |
| NGCCTAN | 18232.3 | 1.32 | |
| NGACAY | 12090.2 | 1.12 | these two motifs combining with GTCAC (conflevel = 3847.4) form GWCAY motif |
| NGTCATN | 12902.7 | 1.33 | |
| NACGTN | 10060.5 | 1.07 | |
| NCCNNGG | 8993.5 | 0.74 | |
| NGAGGN | 8984.4 | 0.28 | high confidence level, moderate signal shift |
| ATTAATN | 10246.2 | 0.96 | |
| NGGWCWAN | 10589.9 | 0.55 | These two motifs form GGWCNA motif |
| GGWCNAN | 9547.2 | 0.46 | |
| NGTACN | 7495.2 | 1.07 | |
| NCGACGN | 6267.5 | 1.02 | these two motifs form CGWCG motif |
| NCGTCGN | 5088.5 | 0.51 | |
| NTCGAN | 4788.8 | 0.44 | Rather low signal shift and confidence level, should be verified manually. |

| | | | |
|---|---|---|---|
| | | | 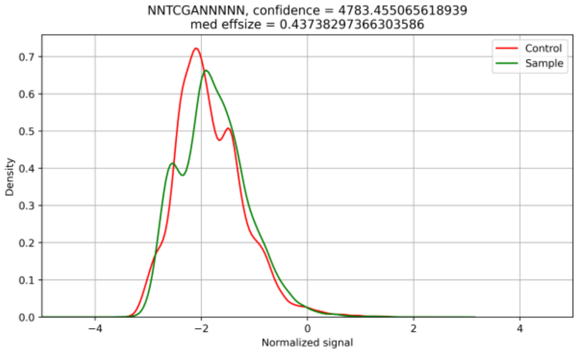<br>Actually, the absence of a common mode indicates that it is an individual motif. |
| CCTAAN | 4838.8 | 0.16 | no significant signal shift (actually it is a cropped submotif of GCCTA) |
| NGGGCTAN | 4508.5 | 0.45 | Rather low signal shift and confidence level, should be verified manually.<br>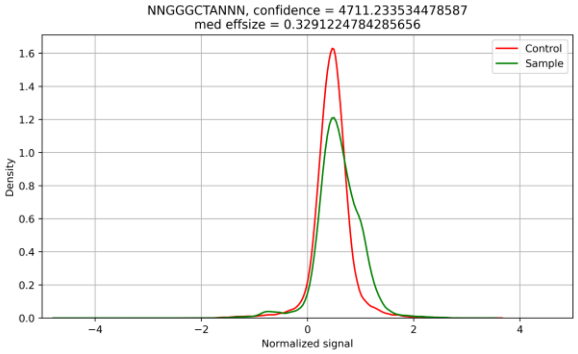<br>The presence of a common mode indicates that it is not an individual motif. |
| NGTCACN | 3847.4 | 1.105 | combining with GTGAC (conflevel = 2307.4) forms GTSAC; combining with GACAY and GTCAT forms GWCAY |
| NTNCCG | 2365.1 | 0.15 | no significant signal shift, low confidence level |
| NGTGACN | 2307.4 | 0.67 | combining with GTCAC (conflevel = 3847.4) forms GTSAC |
| GTCNATN | 2309.9 | 0.19 | no significant signal shift, low confidence level |
| GACNAN | 2326.3 | 0.13 | no significant signal shift, low confidence level |
| NCCNGG | 1977.4 | 0.25 | no significant signal shift, low confidence level |
| GGGCNAN | 1261.5 | 0.17 | no significant signal shift, low confidence level |

confidence level legend:

red <= 3000

3000 < yellow <= 5000

5000 < green

effect size legend:

red <=0.25

0.25 < yellow <= 0.5

0.5 < green

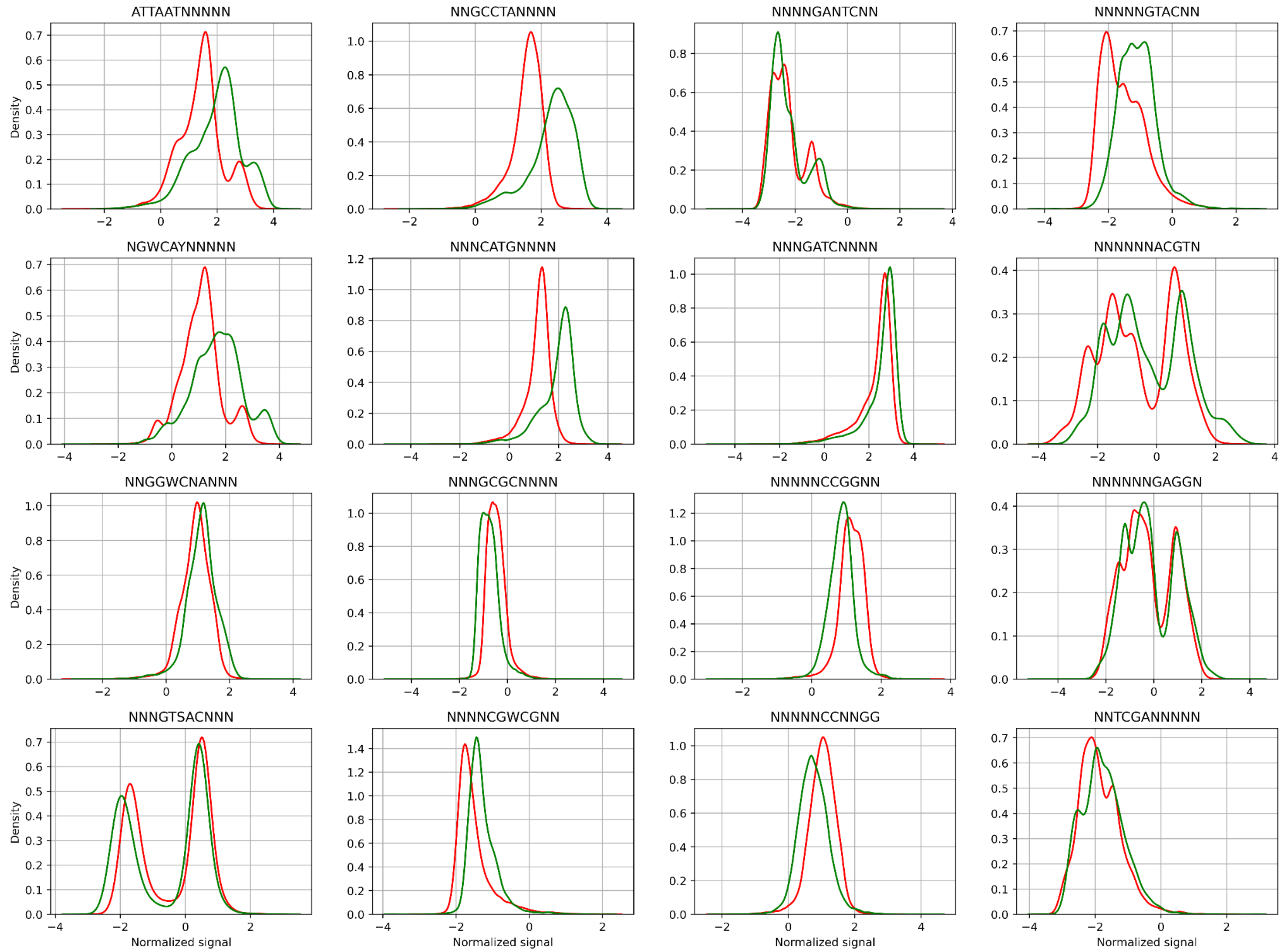

**Supplementary Figure 1.** Signal distributions for methylation motifs discovered in *H. pylori* J99. Red plots represent signal distributions in the WGA sample, green plots - in the native DNA.

### 2. TCNNGA motif in J99 strain

Snapper did not extract the TCNNGA motif as methylated while PacBio did. To verify the results we manually observed all the modified context containing this motif. There were 1255 different 11-mers that had a significant signal shift and contained TCNNGA subsequence. Actually, TCNNGA is a quite highly-represented motif in the J99 genome that might cause a false-positive inference. We checked the presence of other motifs in each context and found that almost all of them (1190 contexts) were explained by the presence of one or more other confirmed methylation motifs (**Supplementary Listing 1**). Among the other 65 probably modified contexts more than half (38) contained homopolymeric fragments which as we found might often cause a false-positive signal shift. Generally, despite a quite high number of TCNNGA-containing contexts the number of unexplained contexts containing only TCNNGA but not any other methylation motifs is insufficient for TCNNGA inference. Thus, we can conclude that TCNNGA is not an individual methylation motif in our J99 strain.

**Supplementary Listing 1.** A shortened list of TCNNGA-contaning contexts that are likely to bring a modified base. The second column shows the presence of a non-TCNNGA motif that explains the signal shift in each case. Only 25 out of 1255 contexts are shown just as an example. In total, 1190 TCNNGA-contaning contexts that have a significant signal change are explained by other motifs.

```
context     actual motif
TCTCGATCCGC GATC
GGAATCAAGAA GANTC
TCAGTCACGAA GTSAC
TTTTCATGACC CATG
ATCATGACTGG CATG
ATCATGAAATT CATG
TAGAGTCCGGA CCGG
GCGTGTCCGGA CCGG
ATCTAGATCCT GATC
TCTTGATCTAT GATC
CACTCATGAGA CATG
TCAAGATCAGA GATC
TACGATCATGA CATG
AAAGATCTAGA GATC
TATCAAGAATC GANTC
GCTTCATGATC CATG
TGCTCATGAAT CATG
ACTCCAGAGGC GAGG
TGCTCATGATA CATG
ATCTCCTGATC GATC
TCAAGATCGCT GATC
TCTTGATCTTG GATC
CACTCATGATC CATG
GATCATGAAAT CATG
GATCATGATTA CATG
```

### 3. A45 extracted motifs

**Supplementary Table 2.** Snapper output for *H. pylori* A45.

| MOTIF | conflevel | effsize | comment |
|---|---|---|---|
| NCATGN | 32951.2 | 1.59 | |
| NGCGCN | 32109.6 | 1.11 | |
| NTGCAN | 19535.3 | 1.00 | |
| NGAACN | 21296.2 | 0.79 | |
| NGGCCN | 22838.8 | 1.42 | |
| NGATCN | 24323.8 | 0.46 | |
| NCCAGN | 22631.8 | 0.72 | |
| NCCATCN | 29952.0 | 1.54 | |
| NGAHTCN | 25034.2 | 0.72 | should be merged with NGANTC to NGANTCN |
| NGGGGAN | 14548.9 | 0.58 | GGAGA and GGGGA form GGRGA |
| ATTAATN | 16256.6 | 1.26 | |
| TCNNGAN | 16322.6 | 0.63 | |
| NTCNGAN | 18074.7 | 0.63 | |
| NGANTC | 11994.4 | 0.37 | |
| NGGAGAN | 11128.4 | 0.60 | GGAGA and GGGGA form GGRGA |
| GTNNACN | 9591.0 | 0.92 | |
| NTCGAN | 4558.2 | 0.62 | |
| CNNGAN | 3066.5 | 0.27 | Rather low signal shift and confidence level, should be verified manually.<br>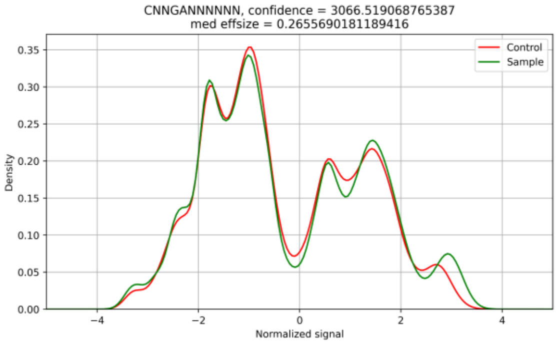<br><br>No noticeable signal shift. Actually, it is a cropped submotif for TCNNGA. |

confidence level legend:
red <= 3000
3000 < yellow <= 5000
5000 < green

effect size legend:
red <=0.25
0.25 < yellow <= 0.5
0.5 < green

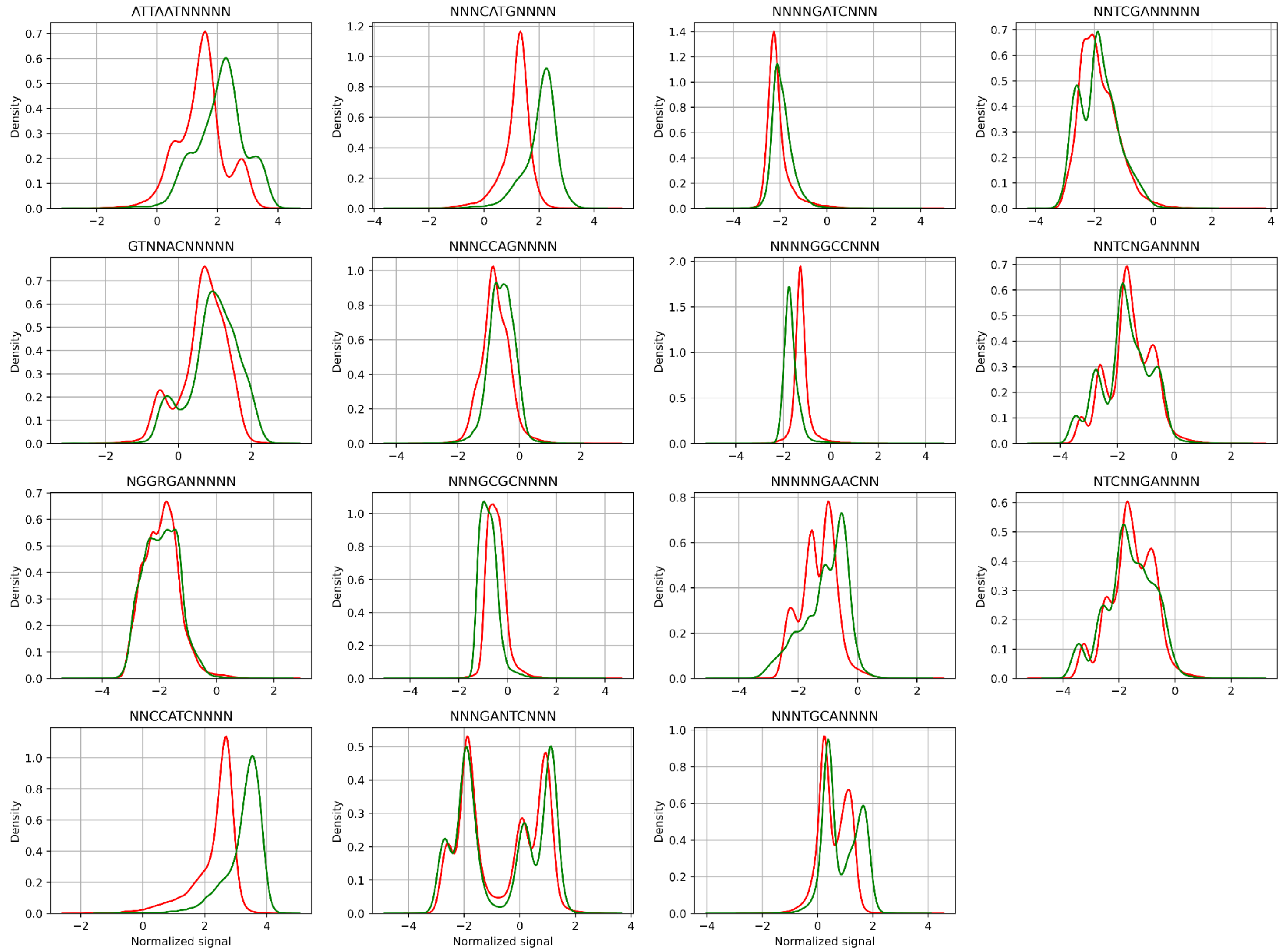

**Supplementary Figure 2.** Signal distributions for methylation motifs discovered in *H. pylori* A45. Red plots represent signal distributions in the WGA sample, green plots - in the native DNA.

### 4. *hpy* mutant analysis

**Supplementary Table 3**. Snapper output for *H. pylori* J99-*hpy* mutant.

| MOTIF | conflevel | effsize | comment |
| --- | --- | --- | --- |
| NCATGN | 70292.8 | 1.54 | |
| NCCAAKN | 68588.4 | 0.52 | |

### 5. *hp1352* mutant analysis

**Supplementary Table 4.** Snapper output for *H. pylori* J99-*hp1352* mutant.

| MOTIF | conflevel | effsize | comment |
| --- | --- | --- | --- |
| NGANTCN | 99349.1 | 0.66 | |
| NGANT | 4009.8 | 0.06 | cropped submotif for NGANTCN |

### 6. *hp91/92* mutant analysis

**Supplementary Table 5.** Snapper output for *H. pylori* J99-*hp91/92* mutant.

| MOTIF | conflevel | effsize | comment |
| --- | --- | --- | --- |
| NGATCN | 68280.2 | 0.69 | |
| NCCAATN | 63263.7 | 1.17 | submotif for NCCAAK |
| NCCAAGN | 83480.3 | 0.53 | submotif for NCCAAK |

### 7. *hp944* mutant analysis

**Supplementary Table 6.** Snapper output for *H. pylori* J99-*bc13* mutant.

| MOTIF | conflevel | effsize | comment |
| --- | --- | --- | --- |
| NCCAGN | 83270.4 | 0.70 | |
| NTCTN | 3174.7 | 0.32 | Rather low signal shift and confidence level, should be verified manually. 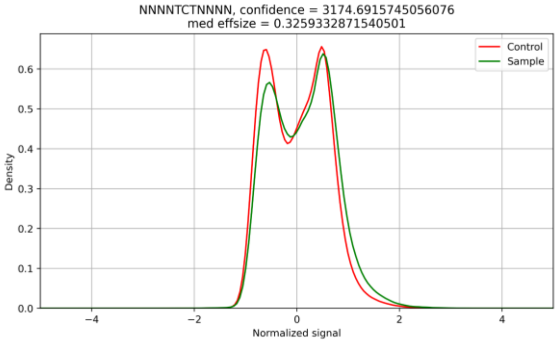  Actually, here we can see a small but noticeable |

| | | | |
|---|---|---|---|
| | | | signal shift for both modes. TCT sequence might be part of an unknown I type R-M system influenced by CCAG-specific MTase deactivation. Snapper is not aimed to infer long methylation sites, so this case should be resolved individually. |

### 8. Signal distribution shifts in deactivated methylation sites in four A45 mutants.

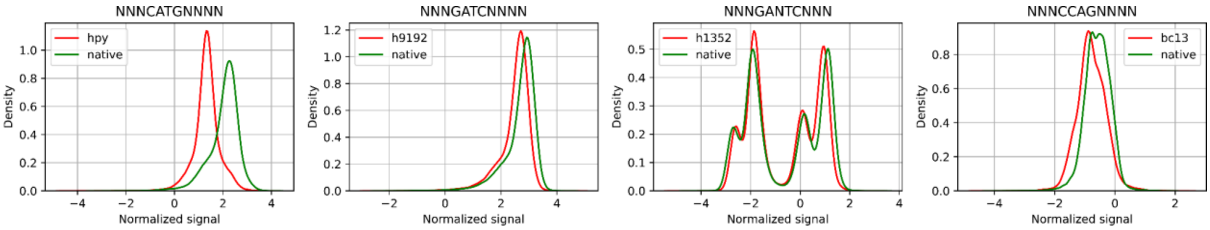

A B C D

**Supplementary Figure 3**. Signal shifts for inactivated methylation sites in *hpy* (A), *hp91/92* (B), *hp1352* (C) and *hp944* (D) mutants. Here, the red plots represent the motif signal distribution in the native DNA sample, green plots - the signal distribution in the corresponding mutant.

### 9. The list of proteins significantly overrepresented in *hpy* and *hp91/92* strains compared with the wild type and the *hp1352* mutant

Non-target proteomic analysis was carried out for wild A45, *hpy*, *hp1352* and *hp91/92*. For each sample, three biological repeats were analyzed, each in 3 technical repeats. In this study the proteomics data were used for identification of a new MTase that turned out to be active only in *hpy* and *hp91/92* mutants but not in the wild type, so, we were interested in the proteins that were significantly overrepresented in *hpy* and *hp91/92* mutants.

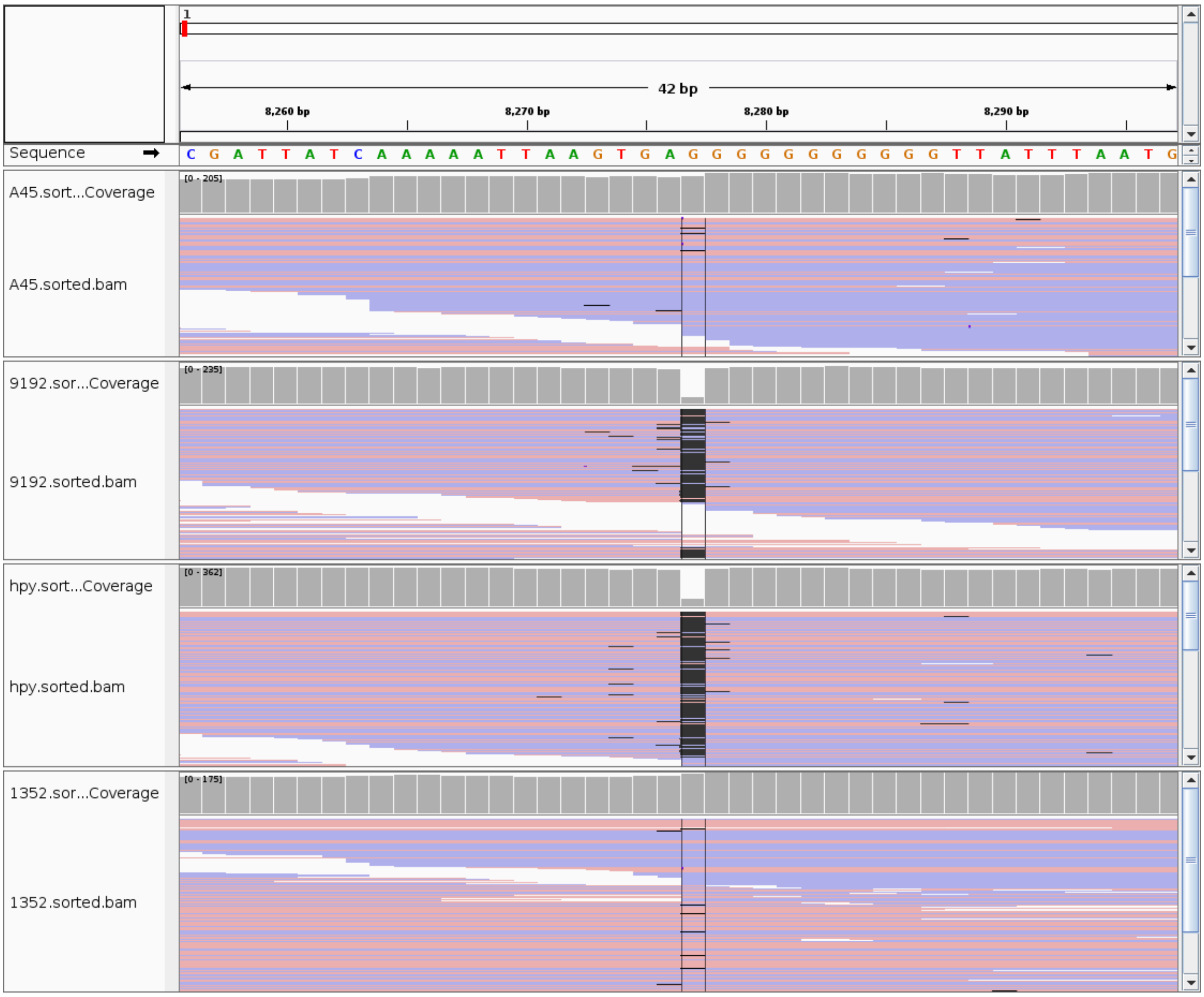

**Supplementary Figure 4**. Mapping Illumina reads to the reference genome sequence wild strain *H. pylori* A45. IGV snapshots with squished coverage mode, pink and blue segments were aligned to the forward and reverse strand, respectively. A black line represents a deletion with the event size.

### 10. Supplementary methods

As we have only one promoter region $P_{Cm}$ from *catGC* cassette, successfully expressing in *H. pylori* cells, we used this promoter in both cases: as a part of chloramphenicol resistant amplicon and fused with promoterless kanamycin resistance gene for kanamycin amplicon. The *catGC* cassette and chloramphenicol resistance gene promoter region $P_{Cm}$ were amplified from the plasmid pHel2 [1] using ACm-F/ACm-R and ACmp-F/ACmp-R primer set, respectively. Promoterless kanamycin resistance gene were amplified from the plasmid pEGFP-N1 (https://www.addgene.org/vector-database/2491/) using AKan-F/AKan-R primer set (Supplementary Table 7). All inner primers were constructed in an overlapping manner for latter two-step PCR amplification of full-length amplicons. (Supplementary Figure 5).

Two-step PCR amplification was realized as follows. On the first step two *H. pylori* flanking fragments and resistance gene fragment in equal concentrations 10 ng were incubated without primers in a final volume of 45 µL containing 1x Tersus plus buffer, 100 µmmol/L deoxynucleoside triphosphate, 0.4 U of Tersus polymerase (Evrogen, Russia) in thermocycler T100 (BioRad, USA) for 8-10 cycles for overlapping fragment parts could «stick» to each other. At the second stage, the appropriate primers, 20 µmmol/L of each primer, were added and mixtures were amplified for 25 cycles (Supplementary Figure 6). In both cases two-step PCR amplification resulted in two full-length fragments 1671 bp for *hp0008* and 1567 bp for *hp0944* gene disruption, respectively.

The amplicon was purified using Cleanup Standard Kit (Evrogen, Russia), cloned into the pCR2.1 vector system using The Original TA Cloning Kit (Invitrogen, USA ) according to the manufacturer's recommendations and transformed into *E. coli* Top10 strain. Cloning was verified by PCR with the M13F/R vector-based primer set (Supplementary Table 7). The plasmids pCR2.1-8, pCR2.1-9 containing the expected amplicon (Supplementary Table 8) were purified using the QIAprep® Spin Miniprep Kit (Qiagen, Germany) and sequenced. Plasmids with verified amplicon nucleotide sequence were digested with restriction enzymes using XbaI/KpnI sites for *hp0008* gene disruption amplicon and KpnI/NotI sites for *hp0944* gene disruption amplicon. Amplicons were purified from agarose gel in appropriate concentrations and used to transform *H. pylori* A45 cells as described earlier.

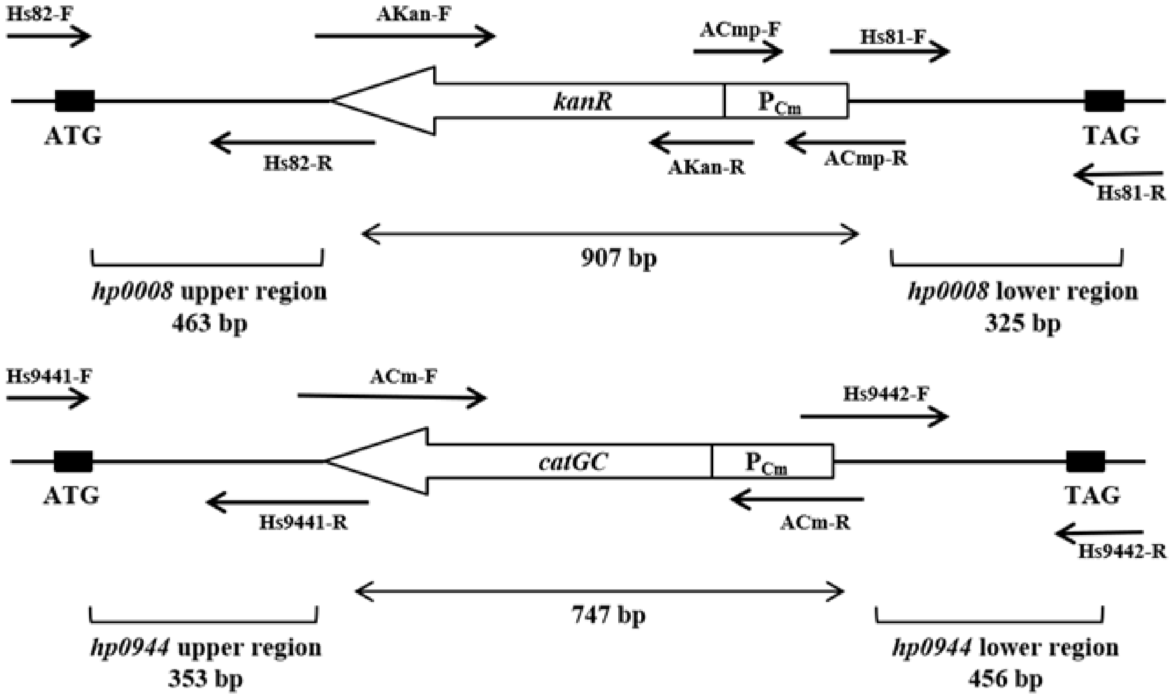

**Supplementary Figure 5.** Schematic map of the primer location and amplicons structure used for hp0008 and hp0944 gene disruption in *H. pylori* A45 strain.

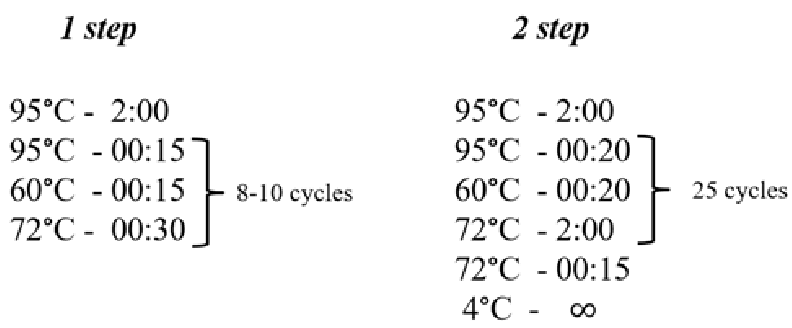

**Supplementary Figure 6.** Two-step PCR conditions used for amplification of full-length fragments 1671 bp for *hp0008* and 1567 bp for *hp0944* gene disruption in *H. pylori* A45 strain, respectively .

**Supplementary Table 7.** Oligonucleotides used in this study.

| Primer set | Sequence | Product |
|---|---|---|
| Hs9441-F<br>Hs9441-R | AATCCACACCCTTTGGATAACT<br>GGGGCGTAACTAATGGAGCGTGGATTAGCA | hp0944 upper region<br>353 bp |
| ACm-F<br>ACm-R | ACGCTCCATTAGTTACGCCCCGCCCTGCCA<br>GTGAGCTTAAACCGCCATATTGTGTTGAAACAC | chloramphenicol resistance cassette |
| Hs9442-F<br>Hs9442-R | ACAATATGGCGGTTTAAGCTCACTTGTCCGTC<br>GGGAGATAGAAGCATAGAAGTT | hp0944 lower region<br>456 bp |
| Hs82-F<br>Hs82-R | TCACGCCGTCTTGTTTGAGC<br>TTTGAGAACTTAAACGAATCCACTAGCAAG | hp0008 upper region<br>463 bp |
| AKan-F<br>AKan-R | GTTTAAGTTCTCAAAATCAGAAGAACTCGT<br>AGCTAAAATGATTGAACAAGATGGATTGCA | promoterless kanamycin resistance gene |
| ACmp-F<br>ACmp-R | TCAATCATTTTAGCTTCCTTAGCTCCTGAA<br>CTGGATACACGCCATATTGTGTTGAAACA | chloramphenicol promoter region |

| | | |
|---|---|---|
| Hs81-F<br>Hs81-R | CAATATGGCGTGTATCCAGCGTGAGCTTATC<br>AGGCTTAGCGACTTTACACCA | hp0008 lower region<br>325 bp |
| 23s-F<br>23s-R | GTAAGCCATAGAAAGTGATAGCCT<br>CTTAACCTTGCCAGATACCACA | primers for verification (complimentary to 23s RNA H. pylori gene) |
| Cms-F<br>Cms-R | ATCCCATATCACCAGCTCAC<br>GGAATTCCGTATGGCAATG | primers for verification (complimentary to chloramphenicol resistance gene) |
| M13f<br>M13r | TGTAAAACGACGGCCAGT<br>CAGGAAACAGCTATGAC | primers for verification (complimentary to pCR2.1 vector) |

**Supplementary Table 8.** Plasmids and strains used in this study.

| **Plasmid/Strain** | **Description** | **References** |
|---|---|---|
| PLASMIDS | | |
| pGEM-HpyIM-κ | pGEM-T-Easy vector system containing full-lenth amplicon for *HpyAI* gene disruption in *H. pylori* A45 strain | [2] |
| pGEM-HpyIIIM | pSK-Kn vector system containing full-lenth amplicon for *hp0091/hp0092* genes disruption in *H. pylori* A45 strain | [2] |
| pSK-HpylVM | pGEM-T-Easy vector system containing full-lenth amplicon for *hp1352* gene disruption in *H. pylori* A45 strain | [2] |
| pCR2.1-8 | pCR2.1 vector system containing full-lenth amplicon for *hp0008* gene disruption in *H. pylori* A45 strain. | This study |
| pCR2.1-9 | pCR2.1 vector system containing full-lenth amplicon for *hp0944* gene disruption in *H. pylori* A45 strain. | This study |

| | | |
|---|---|---|
| *H. PYLORI* STRAINS | | |
| A45 | WT | [2] |
| *hpy* | A45 mutant strain with disrupted in the HpyAI gene by homologous recombination | [2] |
| *hp91/92* | A45 mutant strain with disrupted in the *hp0091/hp0092* genes by homologous recombination | [2] |
| *hp1352* | A45 mutant strain with disrupted in the *hp1352* gene by homologous recombination | [2] |
| A458 (*hp8*) | A45 mutant strain with disrupted in the *hp0008* gene by homologous recombination | This study |
| A459 (*hp944*) | A45 mutant strain with disrupted in the *hp0944* gene by homologous recombination | This study |

### 11. Plasmids used for gene inactivation restriction-modification systems

#### Plasmid Construction pGEM-HpyIM-к (*hpy*)

Two fragments of the gene encoding HpyAI methyltransferase of *H. pylori* strain A45 labeled IMl and IM2 were amplified using primers 1IMfw, 1IMrev, 2IMfvv, 2IMrev (Supplementary Table 9). For the full-length gene assembly, the following reaction mixture was composed: 200 ng of each IM1 and IM2 fragments, 20 pM of each primers 1lmfw and 2Imrev, 2mM dNTPs, 1xPCR buffer, 4u of Taq polymerase. PCR was performed with the following amplification conditions: 94°C – 30 sec, 45°C – 1 min, 72°C – 1 min. 30 sec. A total of 35 amplification cycles were performed. The resulting fragment, after being treated with Xhol restriction enzyme, was cloned into the pGEM-T-Easy plasmid (Promega, USA) which resulted in formation of the plasmid pGEM-HpylM. The *aphA-3* kanamycin resistance gene was amplified from the pHel3 plasmid [1] using primers X-aph-fw and X-aph-rev, then the resulting fragment was digested with restriction enzyme Xhol and cloned into pGEM-HpylM plasmid treated with Xhol restriction enzyme, which resulted in formation of the plasmid pGEM-HpyIM-k.

#### Plasmid Construction pGEM-HpyIIIM (*hp91/92*)

Fragments of the *hp0091* and *hp0092* genes of the A45 *H.pylori* strain labeled 91 and 92 were amplified using primers 91fw, 91revKn, 92fw, 92revKn (Supplementary Table 9). The kanamycin resistance gene (aphA-3) was amplified from the pHel3 plasmid [1] using primers Knfw and Knrev. For cassette assembly the following reaction mix was used: 200 ng of fragments 91, 92 and *aphA-3*; 20 pM of each primers 91fw and 92rev, 2mM dNTPs, 1xPCR buffer, 4u of Taq polymerase. Amplification was performed by 35 cycles of PCR with following conditions: 94°C – 1 min, 45°C – 1 min, and 72°C – 4 min. Assembled cassette then was cloned into pGEM-T-Easy plasmid (Promega, USA) and the plasmid pGEM-HpyIIIM was formed.

#### Plasmid Construction pSK-HpylVM (*hp1352*)

Fragments of the *hp1351* and *hp1352* genes of the A45 *H. pylori* strain labeled 1351 and 1352 were amplified using primers gantcfw1, gantcrev1, gantcfw2, gantcrev2 (Supplementary Table 9). Fragment 1351 was digested by restriction endonucleases HindllI and Xhol, and then cloned into the pSK-Kn plasmid, resulting with the plasmid pSK-1351-Kn. Fragment 1352 was digested by restriction endonucleases PstI and Xbal, and then cloned into the pSK-1351-Kn plasmid, resulting in formation of the plasmid pSK-HpyIVM.

**Supplementary Table 9.** Oligonucleotides used to generate A45 mutant strains *hpy*, *hp91/92* and *hp1352*.

| **name** | **sequence 5'–3'** |
|---|---|
| 1Imfw | TACTCGAGTACTGCCCGCTAAAGCC |
| 1IMrev | CCTAAGCTTGCGAGGCAATACGGGGC |
| 2IMfw | CACTGCAGCCACGCACACCACACATCGC |
| 2IMrev | TTTCTAGAGGGTTCTACGGCTGTAGG |
| X-aph-fw | TCCTCGAGGATCTTTTAGACATCTAAAT |
| X-aph-rev | CGCTCGAGTCGATACTATGTTATACGCC |
| 91fw | GTCTTAAGACAAGCAATAGG |
| 91revKn | GATGTCTAAAAGATCCGCTCCAACCCCGTTTGACG |
| 92fwKn | GATACAATATGCGGCAGCTGCCGTTATTTGACGC |
| 92rev | AGGATCGCCAATGAGAG |
| Knfw | GATCTTTTAGACATCTAAAT |
| Knrev | TCGATACTATGTTATACGCC |
| gantcfw1 | CTCTCGAGGACGCTAATTGCTGATAAATCG |
| gantcrev1 | GAAGCTTTGCGAGACGACCATGACAGCGCC |

| | |
|---|---|
| gantcfw2 | CTCTGCAGCCTTGCGCATCTTTTAGTCTTTCG |
| gantcrev2 | ATCTAGAGAAAACTTAAACACTATCATAG |

**Supplementary Table 10.** The list of proteins significantly overrepresented in *hpy* and *hp91/92*.

| run ID | Protein IDs | | | | | strain |
|---|---|---|---|---|---|---|
| | hp_0008 | hp_0009 | hp_0010 | hp_0927 | hp_0928 | |
| LFQ intensity AO0102_1 | 9810800000 | 7322500000 | 10202000000 | 1190400000 | 1529000000 | *hpy* |
| LFQ intensity AO0102_2 | 9364100000 | 7110000000 | 10412000000 | 1596300000 | 1380800000 | *hpy* |
| LFQ intensity AO0102_3 | 9217600000 | 6737600000 | 10113000000 | 1658200000 | 1688300000 | *hpy* |
| LFQ intensity AO0103_1 | 9362900000 | 6948900000 | 9295500000 | 1165200000 | 970480000 | *hpy* |
| LFQ intensity AO0103_2 | 9665300000 | 6658300000 | 9460100000 | 1132600000 | 998490000 | *hpy* |
| LFQ intensity AO0103_3 | 8736100000 | 6376000000 | 9482000000 | 1128500000 | 1245900000 | *hpy* |
| LFQ intensity AO0104_1 | 9360800000 | 6591000000 | 10903000000 | 1131000000 | 1323500000 | *hpy* |
| LFQ intensity AO0104_2 | 8843400000 | 7027400000 | 10689000000 | 1018700000 | 1583300000 | *hpy* |
| LFQ intensity AO0104_3 | 9332200000 | 6635600000 | 10548000000 | 1356200000 | 1537700000 | *hpy* |
| LFQ intensity AO0111_1 | 5706700000 | 4080700000 | 6088600000 | 375560000 | 669250000 | *hp91/92* |
| LFQ intensity AO0111_2 | 5856900000 | 4130300000 | 5813000000 | 475570000 | 348050000 | *hp91/92* |
| LFQ intensity AO0111_3 | 5860100000 | 3564400000 | 6109200000 | 538440000 | 717000000 | *hp91/92* |
| LFQ intensity AO0112_1 | 10047000000 | 5009000000 | 7978300000 | 645450000 | 494010000 | *hp91/92* |
| LFQ intensity AO0112_2 | 5687900000 | 5605000000 | 7879400000 | 425440000 | 353280000 | *hp91/92* |
| LFQ intensity AO0112_3 | 6209200000 | 3999900000 | 7302800000 | 416250000 | 248620000 | *hp91/92* |
| LFQ intensity AO0113_1 | 6562300000 | 5895700000 | 7702500000 | 201770000 | 158900000 | *hp91/92* |
| LFQ intensity | 0 | 0 | 0 | 0 | 0 | A45 |

| | | | | | | |
|---|---|---|---|---|---|---|
| AO0117_1 | | | | | | |
| LFQ intensity AO0117_2 | 0 | 0 | 0 | 0 | 0 | A45 |
| LFQ intensity AO0117_3 | 0 | 0 | 0 | 0 | 0 | A45 |
| LFQ intensity AO0118_1 | 0 | 0 | 0 | 0 | 0 | A45 |
| LFQ intensity AO0118_2 | 0 | 0 | 0 | 0 | 0 | A45 |
| LFQ intensity AO0118_3 | 0 | 0 | 0 | 0 | 0 | A45 |
| LFQ intensity AO0119_1 | 0 | 0 | 0 | 0 | 0 | A45 |
| LFQ intensity AO0119_2 | 1145400000 | 0 | 0 | 0 | 0 | A45 |
| LFQ intensity AO0119_3 | 0 | 0 | 0 | 0 | 0 | A45 |
| LFQ intensity AO0123_1 | 0 | 0 | 0 | 0 | 0 | *hp1352* |
| LFQ intensity AO0123_2 | 0 | 0 | 0 | 0 | 0 | *hp1352* |
| LFQ intensity AO0124_1 | 0 | 0 | 0 | 0 | 0 | *hp1352* |
| LFQ intensity AO0125_1 | 0 | 0 | 0 | 0 | 0 | *hp1352* |
| LFQ intensity AO0125_2 | 0 | 0 | 0 | 0 | 0 | *hp1352* |
| LFQ intensity AO0125_3 | 0 | 0 | 0 | 0 | 0 | *hp1352* |

**Supplementary Table 11.** BLAST annotation of the proteins overrepresented in *hpy* and *hp9192* mutants.

| gene | protein sequence | annotation |
|---|---|---|
| *hp0008* | MQNKEIGEEKSVKEKNLEVFNRYFPGCLSIENDDKLTLDTG RLKALLGDFSEIKEEGYGLDFVGKKIALNQAFKKNNKILKP LNESTSKHILIKGDNLDALKILKQSYSEKIKMIYIDPPYNTK NDNFIYSDDFSQSNEETLKQLDYSKEKLDYIKNLFGSKCHS GWLSFMYPRLLLAKDLLKQDGVIFISIDDNECAQLKLLCDE IFGEGNFVAEMPRLTKKAGKSTNQIAKNHDYVLCYQKNNI NFKQIDIDENDYPLKDEFYNERGGYKLNQNLDYNSLQYNK KMDYEIVISNEKFYAGGLETYTERQKGNFGTIDWVWRWS KAKFDFGLANGFVEVKNNRIYTKTYTKAKISDSKPYKIEYF NRTKNISSIEFLDNKYSNDMSNKKLQSIFNVKNIFDYSKPVE | site-specific DNA methyltransferase (broken) |

| | | |
|---|---|---|
| | LISFLIDQTTEKGDIILDFFAGSGTTAHAVLESNKSDYQKLSE<br>GGGVI | |
| *hp0009* | MRGGGLFNGLNAAFKERRFILVQLDEKIDPKKNKSAYDFC<br>LNTLKSPSPSIFDITEERIKRAGAKIKEACPHLDVGFRAFEIID<br>DETHANDKNLSQAHQKDLFAYSNPKKRETQTILIKLLGCEG<br>LELTTPINCLIENALYLALNTAFIVGDIEMSEVLENLKDKGV<br>EKISVYMPAISNDRLCLELGSNLLDLKLESGDLKIRG | site-specific DNA methyltransferase (broken) |
| *hp0010* | MKIKFKRLDYQEQCRDQILGVFKGIYLREPENDAQRISNPV<br>FEIGEIKDLLLENIENLRSKQKITQGSVGIDKSLNCDILMETG<br>TGKTFCFLECVYSLHKNYHLSKFIVLVPSNAIKLGVLKSVEI<br>TREFFKSEYSTHLESYEDIRSFILASNHKCCVLVMTFSAFNK<br>EKNTINKSCLENTNLFNGAKSYMQALASISPVVIMDEPHRF<br>LGDKTKKYLEQLNALITLRFGATFKDDYKNLIYALDSKKA<br>FDCALVKSISVASVGESNECFLELKGVVKIQNGYEAMINYT<br>NLENKIQSVKVKKHDNLGALTQISALEDYIVENITKTEARF<br>LNGFNLLLDQKEPFSHLLEGEQEVMLKEAIKSHFEREEGLF<br>KKGIKALCMVFISGVNSYLSENEKPAKLALLFEKLYQQKLE<br>EVLKKEDLDENYRAYLERTKDNIQKVHGGYFAKSKKESDE<br>AQVIALILKEKEKLLSFESDLRFIFSQWALQEGWDNPNVMT<br>ICKLAPSHSHITKLQQIGRGLRLAVNDKGERITKEHADFDFV<br>NELVVIVPQVEGDFVGAIQQEISEHSLIKQVFSGEELEKSGIV<br>KKGYYGALLEKLESLGFGEKTDDENFKLTLNQNEFLEKEP<br>ELEKLKDEKYLNLEKLKGFLKDRLIGNSRVRNKNERKSEKI<br>KINKENFKKFETLWEGLNHQARIAYAIDSESLIDEIVKNIDS<br>SFNVKSKIVSVTTHKKVETMGNNAKTEIFEQKSACVWSLH<br>EFISALSNKVKLSFKSVAKVLENIDENKFDLIKKNEQESLRR<br>LEELFLEIIYQNIRDRISYQMRETTIKDRKNDAFYDEKGEIRE<br>FLDGSLVGDKYEIKNSSTQEKCLYENFMQVESEIEKDTIEES<br>NDTKIIVFGKLPRVKIPIGLNQTYSPDFGYVVENNDKKVLL<br>VVETKGVENESELHEEEKRKISTAKKFFEALKKQGVNIEYK<br>TKIKKDQLSALINEVLNRKD | type III restriction-modification system endonuclease |
| *hp0927* | MLLEQIQAHSSGFEKFIVKTLGIQNVENFINNWYGKQSLSSF<br>ANNFVPGGLNQALDKIGSSTDAKDLQSFLDKTTFGDILNQ<br>MINQAPLINKLISWLGPQDLSVLVNIALNSITNPSKELTSTIS<br>SIGEKALNDLLGEGVVNKIMSNQVLGQMINKIIADKGFGGV<br>YHQGLGSILPKSLQKELEQFGLGSLLGSRGLHNLWQKGNF<br>NFLAKDYVFVNNSSFSNATGGELNFVAGKSIIFNGKNTINFT<br>QYQGKLSFISKDFSNISLDTLNATNGLILNAPRNDISVQKGQ<br>ICVNVLNCMSEKKTNPSTSSAPTDETLEVNANNFAFLGTIK<br>ANGLVDFSKVLQNTTIGTLDLGANATFKANNLIVNNAFNN<br>NSNYRVNISGNLNVVKGAALSTNENGLNVGGDFKSEGSLIF<br>NLNNPTHQTIINVTGASTIMSYNNQTLINLNTQLKQGSYTL<br>MDAKRMLYGYDNQIIRQGSLSDYLKLYTLIDFNGKRMQLN<br>GDSLSYDNQPVNIKDGGLVVSFKDNQGQMVYSSILYDKVQ<br>VSVSDKPMDIHAPSLEYYIQRIQGSGGLNAIKSAGNNSIMW<br>LNELFVAKGGNPLFAPYYLQDNPTEHIVTLMKDITSALGML<br>SNSNLKNNSTDALQLNTYTQQMSRLAKLSNFASFDSTDFSE<br>RLSSLKNQRFADAIPNAMDVILKYSQRDKLKNNLWATGVG<br>GVSFVENGTGTLYGVNVGYDRFIKGVIVGGYAAYGYSGFY<br>ERITNSKSDNVDVGLYARAFIKKSELTFSVNETWGANKTQI<br>NSNDTLLSMINQSYKYSTWTTNAKVNYGYDFMFKNKSIIL | vacuolating cytotoxin domain-containing protein |

| | | |
|---|---|---|
| | KPQIGLRYYYIGMTGLEGVMNNALYNQFKANADPSKKSV<br>LTIDLALENRHYFNTNSYFYAIGGVGRDLLVRSMGDKLVR<br>FIGDNTLSYRKGELYNTFASITTGGEVRLFKSFYANAGVGA<br>RFGLDYKMINITGNIGMRLAF | |
| *hp0928* | MTYRNSKIDLKNERFSKNRSFKGVKKKIAKKHKAKNLSLI<br>AHAFKTQSNLSASFNKKIFLGLGFVSALSAEDYKSSVYWLN<br>SVNENNSHKSYYVSPLRTWAGGSRSFTQNYNNSQLYIGTK<br>NASATPNNSSIWFGEKGYVGFITGVFKAKDIFITGAVGSGN<br>EWKTGGGAILVFESSNELNANGAYFQNNRAGTQTSWINLIS<br>NNSVNLTNTDFGNQTPNGGFNAMGRKITYNGGIVNGGNFG<br>FDNVDSNGTTTISGVTFNNNGALTYKGGNGIGGSITFTNSNI<br>NHYKLNLNANSVTFNNSTLGSMPNGNANTIGNAYILNANN<br>ITFNNLTFNGGWFVFNRPDANVNFQGTTTINNPTSPFVNMT<br>GKVTINPNAIFNIQNYTPSIGSAYTLFSMKNGNITYNNVNNL<br>WNIIRLKNTQATKDNSENATSNNNTHTYYVTYNLGGTLYH<br>FRQIFSPDSIVLQSVYYGANNIYYTNSVNIHDNVFNLKNIND<br>DRADTIFYLNGLNTWNYTNARFTQTYGGKNSALVFNATTP<br>WANGAIPKSNSTVRFGGYEGVNWGKTGYITGTFTADRVYI<br>TGNMMSGNGAQTGGGATLNFVGATEINIAGADFKNLKTTS<br>QNSYMTFIALGDSSGSGKINVSQSDFYDWTGGGYDFTGNS<br>TFDSVNFNKAYYKFQGAENSYTFKNTNFLAGNFKFQGKTT<br>IEKSVLDDASYSFDGINNAFNEDKFNGGSFNFNAKQVNFSG<br>NSFNGGVFNFNNTPKVSFTDDTFNVNNQFKINGAQTTFTFN<br>KGVIFNMQGLLSSLSVGTTYQLLNAKSVDYKDNNNALYQ<br>MLHWISGENPSGKLVDENKTAPSSAKIYNVHFTDNGLTYYI<br>KENFNNGITLTRLCTLGYTHCVNIHNEVFHLKNINNNASNT<br>VFYLNGMTTWKIAGTGVFTQDYSGANSVLVFNQTTPFLAG<br>ANPTSNSVVSFGKTSGAEWGLVGYIQGVFKANQIDITGTIR<br>SGNGAQTGGGATLVFNAEKRLNIANAHLNNDKAGLQDSW<br>MNFIVNNGNLNATNANFSNQTPHGGFNLKANDITFNGGSV<br>SGGGNFGVDNANANGNAVIKNVNFSDNGTLIYKGGENSA<br>GNSLTLENNTFNSYNINARVQNLIFNNNSFNGGSYSFNDTK<br>NTTFKGTNTLINSDPFSRLQGSIAIDNNSIFNIERDLTDNTTY<br>TLLSGNNIKYNNEILADNAFSKNLWNLIHYGGEQGTLLRA<br>DNNTFFVQFTQSNGQKFVFEETFNSSSITYKYFTIHSSLFHT<br>DNDSKDIWSQVRKQFDFIPGKTPVCVGVCYIAPYKNQDLIG<br>SSAFAWSLNFGVTVVGTLLLGSAQEKANDNGGSIWFGKNN<br>LLYLHGNFNATNIFLTNNFNVGNPNAGGGATISFNADETLS<br>ADGLNYTNFQTVAMGLQTSTSQHSWANFNSRLSMEIKNSN<br>FRDFTWGGFNFNSGRIAFENTTFSGWTNINGATESGSSYVN<br>MVANTDLIFTDSILGGGIRYDLKANNIIFNNSQIVIDVSKNV<br>NQSSLNGNVTFNHSRLSVKPNAAINIGDSQTQTTLENASSLS<br>FYNNSVANFNGTTAFNGVSYLNLNPNAQVSFNQANFNNA<br>NVTFYGIPLFGKTPDFGNSARLINFKGNTNFNQATLNLRAK<br>NIHINFQGASTFENNSTMNLAESSQASFNALSVEGETNFNL<br>NNSSLLNFNGNSVFNAPVSFYANNSQISFTKLATFNADASF<br>DLSNNSTLNFQSVLLNGTLNLLGNGANALAINASGNFSFGT<br>QGVLNLSNVNLFDAKNKPLVYNILQAQNIQGLMGNNGYE<br>KIRFYGIQIDKADYSFNNGVHSWSFTNPLNTTETITETLHNN<br>RLKVQISQNGVSNNEMFNLAPSLYFYQKNPYNESSNSYNY<br>TSDKAGTYYLSSNIKSFNQNNKTPGTYNVQNQPLQALHIY<br>NQAITKQDLNMIASLGKEFLPKIANLLSSGALDNLNLNSPN<br>GFETLFGIFEKYGITLNQENWKSLLKIINNFSNTANYDFSQG | vacuolating cytotoxin domain-containing protein |

| | |
|---|---|
| | NLVVGAIKEGQTNTNSVVWFGGEGYKEPCAVGDNTCQMF<br>RQTNLGQLLNSSAPYLGYINANFKAKNIYITGTIGSGNAWG<br>SGGSANVSFESGTNLVLNQANIDAQGTDKIFSYLGQGGIEK<br>LFGEKGLGNALSNIIYEESLNDNAIPKDLANMIPKDLGSKTL<br>SSLLSPTEVNNLLGVSAFKNAIMEILNSKTVGDVFGENGLL<br>NALIL |

### 12. Motif enrichment algorithm.

The input data required by the algorithm are a list of 11-mers that are likely to bring a modified base in the FASTA format and a reference genome in the FASTA format. The optional algorithm parameters are the desired confidence level threshold ***conflevel*** (default is 1000) and the maximum number of extracted motifs ***max_motifs*** (default is 20).

1) The first stage is the generation of ***motif_variants***, the list of all potential motif variants. An individual motif variant is a vector with size of 11 looking as follows:

$$(b_1, b_2, b_3, \dots , b_9, b_{10}, b_{11})$$

where the i-th element represents the nitrogenous base (A, G, T, or C) located in the i-th position of 11-mer.
The maximum size of ***motif_variants*** is limited by the total set of 11-mers presented in the reference genome. Since the algorithm aims to extract short methylation sites we use special heuristics to filter the initial ***motif_variants*** list. Firstly, the number of canonical bases in one particular motif variant is limited from 3 to 6 (that can be extended to 8 during the motif adjustment stage) according to published data. Secondly, the presence of either ‘A’ or ‘C’ base in each motif variant is required since for bacteria only these two bases have been described as methylation positions. (Formally, these constraints are not necessary but they significantly reduce the number of potential motif variants. We tried to implement a total brute force approach but it turned out to be dramatically more time-consuming ).

2) Next, the algorithm generates the ***reference_set*** list containing all 11-mers presented in the reference genome. It will be used as a control set. The set of all 11-mers that are most likely to bring a modified base will be named ***seq_set***.

3) Next, the algorithm performs the chi-square test for each motif presented in ***seq_set*** in order to compare its frequency in ***seq_set*** with the frequency in ***reference_set***. After all motifs in ***seq_set*** are processed, the ***motif_variants*** list is sorted according to the chi-square statistic value in descending order. The top motif is extracted and adjusted.

4) The adjustment process works as follows. All `N` bases in the considered motif variant that are neighboring with canonical bases are iteratively changed to ‘A’, ‘C’, ‘G’, and ‘T’ and the frequency of resulting motifs is compared with the reference. If the A:C:G:T ratio in one particular position in ***seq_set*** is close to ***reference_set***, this base remains to be ‘N’. Otherwise, it changed to a more suitable letter (Y, W, S, H, etc). For the adjustment, the algorithm uses only non-filtered ***seq_set*** regardless of iteration.

5) Next, the algorithm checks if the adjusted motif is a submotif for one of the extracted earlier motifs. If it is, the motif is transformed to its supermotif. For example, if we have already extracted the motif 'NNNCATGNNNN' with higher confidence level, and the current adjusted motif variant is 'NNNNCATGWNN', it will be transformed to 'NNNNCATGNNN'.

6) Thereafter, the algorithm extracts the adjusted motif variant and performs filtering of the ***seq_set*** by removing all 11-mers that match this motif variant.

7) If the confidence level of the last extracted motif is less than ***conflevel*** or the number of extracted motifs equals ***max_motifs***, the algorithm stops. Otherwise, the algorithm goes to 4) and repeats processing of the filtered ***seq_set*** to try to extract the next motif.

The algorithm output is the list of extracted motifs.
